# Indian Wellderly exomes underscore *NRAP* roles in elderly hypertrophic cardiomyopathy

**DOI:** 10.1101/789065

**Authors:** Prasanth Chimata, Sharma Ankit, Radhika Agrawal, Vinay J Rao, Rakesh Koranchery, Ranjith Rajendran, Kurukkanparampil Sreedharan Mohanan, Jayaprakash Shenthar, Sholeh Bazrafshan, Sakthivel Sadayappan, Hisham Ahamed, Perundurai S Dhandapany

## Abstract

Hypertrophic cardiomyopathy (HCM) is a genetic disorder that affects people of all ages, with the elderly population being inadequately studied. It is primarily caused by gene variants that encode proteins involved in the structure and function of the heart muscle. The identification of genes associated with elderly HCM requires ethnic-specific genomic sequences from Wellderly individuals. Currently, no Indian Wellderly dataset is available. To address this, we collected and sequenced the Indian Wellderly population. We built a novel Indian database of healthy aging nucleotide sequences (named i-DHANS) and is newly accessible at <u>IndiCardiome</u>. Utilizing this database and Indian HCM cohort, we identified nebulin-related anchoring protein (NRAP) as a gene associated with elderly HCM. NRAP is crucial for the assembly of myofibrils and transmission of force from the sarcomere to the extracellular matrix. Our functional analysis showed that the identified NRAP variant had significantly reduced interactions with its interacting partners, such as Kelch-like protein 41 (KLHL41) and α-actinin, implying a loss of function. In summary, our findings indicate that *NRAP* is a new elderly cardiomyopathy gene, and our Indian Wellderly database is a valuable resource for identifying ethnic-specific genes for various diseases.

## 1. Introduction

Hypertrophic cardiomyopathy (HCM) affects the heart muscle, causing abnormal muscle thickening and ultimately leading to systolic dysfunction. The clinical manifestations of HCM typically begin in early adulthood (approximately 20 years of age) and progress rapidly in middle-aged individuals (approximately 40 years of age). However, in elderly patients, symptoms may remain silent until the age of 55 years (Lewis & Maron, 1989). Currently, little is known about the genetic basis of elderly and late-onset HCM, and only a few sarcomeric genes, such as myosin-binding protein C3 (*MYBPC3*), cardiac troponin T (*TNNT2*), cardiac troponin I (*TNNI1* and *TNNI3*), myosin light chain 3 (*MYL3*), alpha-cardiac myosin heavy chain 6 (*MYH6*), and beta-cardiac myosin heavy chain 7 (*MYH7*), are associated with this condition(Anan et al., 2007; Niimura et al., 2002). Additionally, exome sequences from individuals above 80 years without any chronic ailments or medications (termed as wellderly individuals) across the globe are also scanty to identify causative gene variants for various diseases, including elderly HCM (Erikson et al., 2016).

In this study, we collected and sequenced exomes from 94 Indian Wellderly individuals. Furthermore, we generated the first Indian Wellderly database ([IndiCardiome](https://instem.res.in/IndiCardiome/)) to discover new genes associated with elderly HCM. By utilizing this database and Indian HCM cohort, we identified a missense mutation in *NRAP* through exome and targeted sequencing. NRAP is a multi-domain scaffolding protein involved in the assembly of cardiac muscle thin filaments, organization of myofibrils, and actin cytoskeleton(Carroll et al., 2001). We conducted *in vitro* studies to examine the functional consequences of this missense mutation, which revealed a loss of function due to reduced interaction with its binding partners, such as KLHL41 and α-actinin, supporting a pathogenic role for this gene in elderly HCM.

## 2. Materials and methods

### 2.1 Selection criteria for Indian Wellderly individuals

We screened various Indian control samples as shown in Figure 1. From these, we selected healthy aging individuals who were defined as individuals aged >80 years with no chronic diseases and were not taking any chronic medications. Individuals with any of the following disease phenotypes that were known to cause frequent and leading old age-related deaths were excluded from enrollment: a) Chronic Infection and autoimmune disease, (b) cardiovascular disease, (c) hypertension, (d) diabetes mellitus, (e) cancer, and (f) brain-related disorders (self-reported). After applying these stringent criteria, we obtained 94 healthy aging individuals and performed whole exome sequencing and analysis using in-house pipelines, as described in section 2.4. Genomic DNA was extracted from the peripheral blood samples of the Wellderly individuals using standard procedures (phenol-chloroform method) for further analysis (Dhandapany et al., 2009).

**Figure 1:**
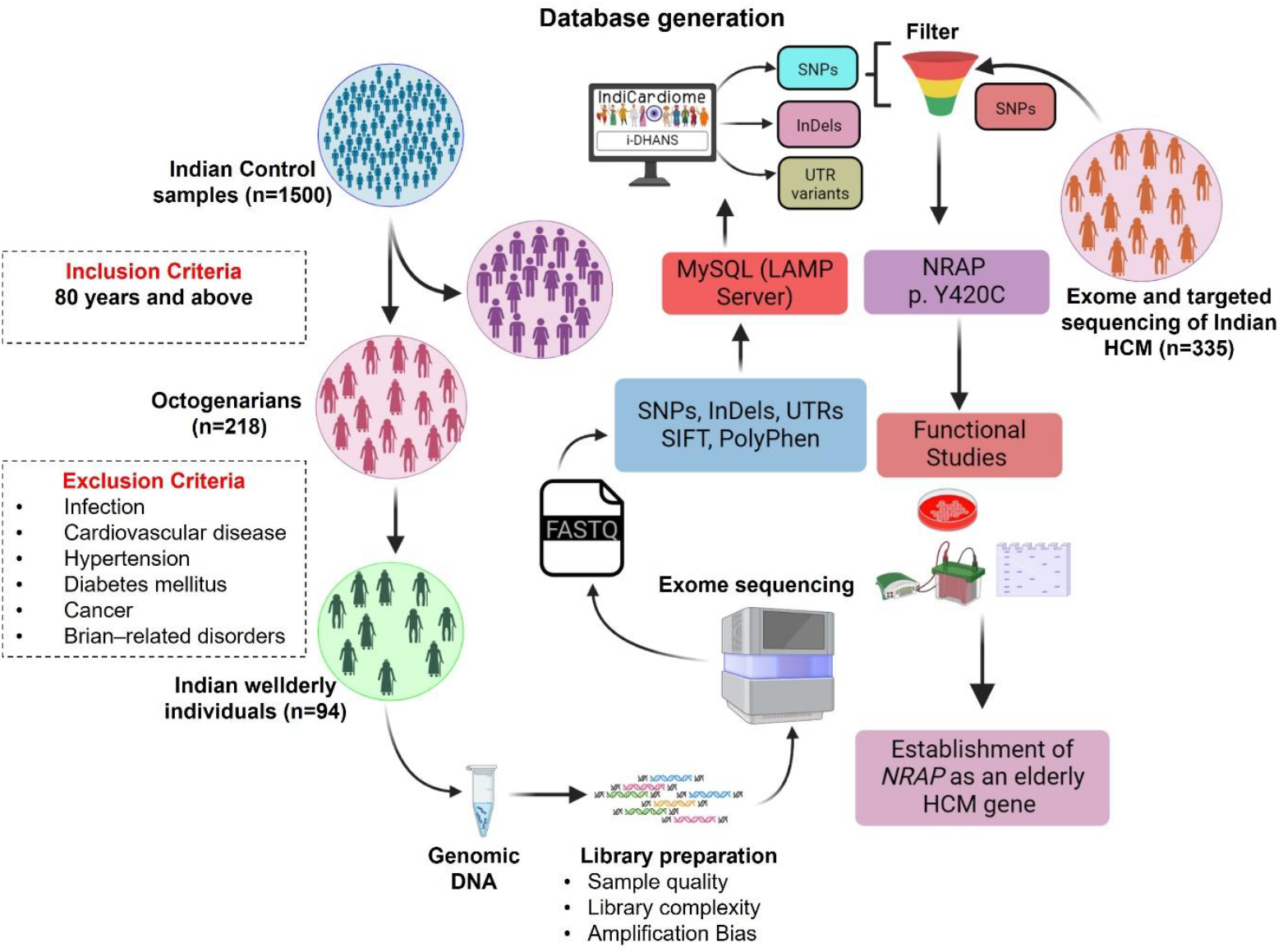
A Schematic representation of study design.

### 2.2 Generation of an Indian database of healthy aging nucleotide sequences (i-DHANS)

#### Data storage and web interface

Genetic variants identified by exome sequencing and analysis of Wellderly individuals were stored and managed using the MariaDB database. Different datasets including missense, insertions/deletions (InDels), truncations, and regulatory region (UTRs) variants are organized in several tables, with each table having a primary key called ‘Node ID’ and a unique identifier called ‘Gene Name.’ To provide a user-friendly web interface for accessing information, the front end was created using Drupal 10 CMS and PHP 8.2.18. The LAMP server platform was used to manage all web services of the i-DHANS and was successfully tested over different operating systems, including Linux, Windows, and the MAC OS SIERRA. These were made publicly accessible by [the IndiCardiome](https://instem.res.in/IndiCardiome/).

#### Data organization and integration

All annotated and filtered variants were stored in the MariaDB database using the LAMP server. A database was created for four different types of datasets, including missense, insertions/deletions (InDels), truncations, and regulatory region (UTRs) variants. The view is linked by multiple tables of different content types, and records are uniquely identified using the field “Gene_Name.” Data can be accessed using gene names. We also string the system IP address of the corresponding user into the backend table of our database for security purposes.

On the homepage of the i-DHANS database, a user-friendly search option is provided for specific gene queries for missense, insertions/deletions (InDels), truncations, and regulatory region (UTRs) variants. Additional information on longevity-associated genetic variations in different ethnicities (including healthy aging Indians) can be obtained. These were made publicly accessible by [the IndiCardiome](https://instem.res.in/IndiCardiome/).

### 2.3 Sampling and diagnostic criteria for HCM patients and controls

Blood samples from HCM patients and their clinical reports were obtained from the Government Medical College, Kozhikode, Sri Jayadeva Institute of Cardiovascular Sciences and Research, Bangalore, and Amrita Institute of Medical Sciences, Kochi. The study was conducted in accordance with the ‘Declaration of Helsinki’ and approved by the respective hospitals and institutional ethics committees. Informed written consent was obtained from all individual participants included in the study.

A standard international protocol was employed to diagnose patients with HCM, which included Probands 1 and 2, as previously described (Dhandapany et al., 2021).The baseline characteristics of the HCM patients for the replication cohort (n=335) are provided in Supplementary Table 1. The control groups consisted of a total of 1029 healthy Indian genomes(A. Jain et al., 2021), 94 Wellderly Indians aged 80 or older (from the i-DHANS database), and 173 Indian Americans. Genomic DNA was extracted from peripheral blood samples of the study participants as described in section 2.1.

### 2.4 Whole exome sequencing

Exome sequencing, analysis, and variant calling were carried out utilizing a custom in-house pipeline, as previously documented in Jain et al. (2022)(P. K. Jain et al., 2022). In brief, DNA libraries were prepared and mixed with capture probes for the targeted regions following the protocol provided by the SureSelect V5 Target Enrichment kit (Agilent Genomics)(Xu et al., 2015). The enriched libraries were amplified using polymerase chain reaction (PCR). Sequencing and targeted re-sequencing were performed by the respective service providers, including the institutional sequencing facility, after de-identifying the patient details. The exomes were sequenced using paired-end 100 base pair reads (100X coverage), and 6 Gb of data was obtained. The sequencing-derived raw image files were processed using BGISEQ-500 basecalling software with default parameters, generating the sequence data as paired-end reads in the FASTQ format. The quality of the raw data was checked using FastQC software, and only qualified reads were used for further processing. Low-quality reads were filtered, and adapters were removed using Trimmomatic.

Exomes were initially aligned onto the human reference genome (GRCh38) using the Burrows-Wheeler aligner version 0.7 (BWA-MEM). The resulting SAM reads (sequence alignment/map format) were transformed into binary mapped format (BAM) via the Samtools software, which is utilized to view the index of binary-aligned sequences. Indexing of mapped reads was performed using the Picard tool package (https://broadinstitute.github.io/picard/). Variant calling was executed using the HaplotypeCaller from the Genome Analysis Tool Kit (GATKv.3.4) (https://software.broadinstitute.org/gatk/). Variants were annotated with the web interface of Annotate Variation (ANNOVAR) software (https://wannovar.wglab.org/). All protein-coding region genomic variations, such as single nucleotide polymorphisms (SNPs), insertion/deletion (InDels), and other known functional elements, such as untranslated regions (UTRs), were detected by HaplotypeCaller of GATK (McKenna et al., 2010). All variant calling files (VCF) for each sample were annotated using wANNOVAR (Wang et al., 2010).

### 2.5 Prioritizing variants in elderly HCM

To determine the potential causal genetic variants in the elderly HCM patient, exome sequencing was carried out on Proband 1 (2II). The following parameters were taken into account for prioritizing variants: (a) coding regions; (b) novel missense; (c) ultra-rare variants present less than 0.01% frequency (Minor Allele Frequency (MAF) ≤ 0.01%) in the reference populations, including the Single Nucleotide Polymorphism database (dbSNP-version 142), 1000 Genomes Project, Exome Sequencing Project (ESP), Korean exomes database (KOVA), Exome Aggregation Consortium (ExAC), Genome Aggregation Consortium (gnomAD) cohort alleles, Genome Asia 100 K, and South Asian (Indian) healthy controls (A. Jain et al., 2021) and the Indian Wellderly Database ([i-DHANS](https://instem.res.in/IndiCardiome/)). Amino acid conservation across species was analyzed by comparing the protein sequences of various vertebrate species using ClustalW2 software. (d) Deleterious variants affecting potential protein functions were identified based on protein damage prediction software using wANNOVAR for Polymorphism Phenotyping v2 (PolyPhen2), Sorting Intolerant From Tolerant (SIFT), and Mendelian Clinically Applicable Pathogenicity (M-CAP), MutationTaster, and Combined Annotation Dependent Depletion (CADD) methods. Furthermore, cardiovascular-related genes were filtered out using the International Mouse Phenotyping Consortium (IMPC) (www.mousephenotype.org/) and Mouse Genome Informatics (MGI) (www.informatics.jax.org/) databases. Finally, a comprehensive literature survey was performed to understand the role of the particular genes in HCM.

### 2.6 Targeted re-analysis of *NRAP* variant in 335 HCM patients

To replicate the association of the *NRAP* gene variant, we performed targeted resequencing in an independent cohort comprising 335 HCM patients. The baseline characteristics of the 335 HCM patients are outlined in Supplementary Table 1. For this purpose, exons and flanking intronic boundaries of *NRAP* were amplified from genomic DNA. Amplified PCR products were gel-isolated with a QiaxII Gel Extraction Kit (Qiagen), sequenced on an ABI3730 DNA analyzer (Perkin-Elmer Corp., Applied Biosystems, Hitachi, Japan) using a BigDye terminator kit (Applied Biosystems), and analyzed on an ABI3730 DNA Analyzer.

NRAP genomic primer forward: GAAAATTTTCAGGGATCTTACCTAG

NRAP genomic primer reverse: CTGTCTTACTCCAACTTCTTGTG

### 2.7 Segregation analysis in the patient’s family members

We performed segregation analysis on five subjects from an HCM family of Proband 1 using Sanger sequencing.

### 2.8 Protein-protein interaction (PPI) analysis

HCM-associated genes were extracted from an earlier published report (Xu et al., 2015), and a protein-protein interaction network was constructed with the identified potential gene (*NRAP*) from the patient’s exome data. PPI network analysis using the STRING database (based on protein co-expression, experimentally determined interaction, and various database annotations) and pathway analysis were performed using ClugoVs from Cytoscape 3.6.0 (www.cytoscape.org/). Only significant pathways with ≤0.05 were considered for pathway enrichment. Conservation alignment was performed using Clustal Omega (www.ebi.ac.uk/Tools/msa/clustalo/).

### 2.9 Plasmid Constructs

The plasmid constructs were generated using the Gibson assembly cloning method for NRAP, KLHL41, and α-actinin from respective cDNA using high-fidelity Q5 polymerase (New England Biolabs). The plasmids were cloned into the respective vectors (pHACE and pcDNA3.1 EGFP) using the primers listed below.

KLHL41 Forward: actctcggcatggacgagctgtacaagatggattcccagcgggag

KLHL41 Reverse: acactatagaatagggccctctagattatagtttagacagtttaaacagatttaagcg

α-actinin Forward: actctcggcatggacgagctgtacatgaaccagatagagcccg

α-actinin Reverse: acactatagaatagggccctctagactacagatcgctctccccgtag

NRAP Forward: ggatcctcgaggccaccatggaattcatgacaaaggactctgtagacc

NRAP Reverse: ttaggcgtagtcaggcacgtcgtaaggatagaattccttatattcaacctcgctgctc

The PCR conditions for the primers listed above were as follows: denaturation at 95°C for 30s, annealing at 56°C for 30s, elongation at 72°C for 2 min for 40 cycles. Site-Directed mutagenesis was carried out for generating p.Y420C in NRAP expressing plasmid (QuikChange Site-Directed Mutagenesis Kit, Stratagene).

### 2.10 Immunoprecipitation (IP) and immunoblotting

HEK293T cells were cultured in Dulbecco’s modified Eagle’s medium (DMEM) supplemented with 10% FBS. The NRAP WT and NRAP p.Y420C constructs were overexpressed along with an equal amount (4 μg) of KLHL41 or α-actinin in HEK293T cells using Jetprime polyplus, following the manufacturer’s instructions in 100 mm dishes. After 24 hours post-transfection, cells were washed with PBS and lysed for 30 min in IP buffer. They were then quantified using the BCA protein assay kit (Pierce™ BCA Protein Assay Kit, Thermo Fisher).

The protein samples were mixed with 6x Laemmli buffer containing 5% β-mercaptoethanol. The samples were incubated at 95°C for 10 min, followed by SDS-PAGE electrophoresis was carried out. The proteins were then transferred onto a PVDF membrane using a Bio-Rad transblot cell. All immunoblot (IB) antibodies were diluted to a concentration of 1:1000 in 3%bovine serum albumin (BSA; HiMedia) in TBST buffer. The blots were blocked for an hour, after which they were incubated with primary antibodies overnight at 4°C, followed by an hour-long incubation with the respective secondary antibodies at room temperature. The process was then completed by developing blots using chemiluminescence on a Westar Antares system (Cyanagen).

For immunoprecipitation (IP), samples containing 250 µg of protein were incubated with 5 μL of their respective primary antibodies overnight at 4°C. The samples were then incubated with 100 µL of recombinant protein G-Sepharose ™ 4B (101242, Thermo Fisher) overnight at 4°C and subsequently washed four times with IP buffer. The protein complexes were eluted with 6x Laemmli buffer and analyzed by SDS-PAGE, followed by western blotting, as described above. The following antibodies were used: anti-HA (IP & IB) (1:1000, 26183, Thermo Fisher), anti-GFP (IP) (MA5-15256, Thermo Fisher), anti-GFP (IB) (1:1000, 2956T, Cell Signaling Technology), and mAb IgG1 Isotype Control (#5415n, Cell Signaling Technology).

### 2.11 Statistical analysis

Statistical significance was determined by using Student’s t-test, with a confidence interval of 95%. A value of p<0.05 was considered statistically significant.

## 3. Results

### 3.1 Generation of i-DHANS, an Indian Wellderly exomes database

To identify genes associated with elderly cardiomyopathy, we generated the first Indian Wellderly Exome Database. For the database, we collected and exome sequenced 94 healthy aging octogenarian Indians who did not have any chronic medical conditions, such as chronic infection and autoimmune disease, cardiovascular disease, hypertension, diabetes mellitus, cancer, and brain-related disorders. These exome datasets were compiled into the Indian Database of Healthy Aging Nucleotide Sequences (i-DHANS) (Figure 1). i-DHANS provides comprehensive information on different types of genetic variation, including missense, insertions/deletions (InDels), truncations, and regulatory region (UTRs) variants in healthy aging elderly Indians. These were made publicly accessible by [the IndiCardiome](https://instem.res.in/IndiCardiome/). Furthermore, it will be helpful in filtering and validating the pathological mutations for any disease, including HCM. For example, we analyzed HCM-associated variants of uncertain significance (VUS) or conflicting interpretation-associated variants from ClinVar/gnomAD and validated them for their disease-causing nature (Supplementary Table 2).

### 3.2 Identification of *NRAP* as an elderly HCM gene using i-DHANS database

Next, we identified a heterozygous missense variant in Nebulin-Related-Anchoring Protein (*NRAP*) (NM_001261463, c.1259A>G, p.Y420C) in exome sequencing and analysis of an affected family with an elderly HCM patient (Proband 1(2II), Figure 2A) using a filtering pipeline (P. K. Jain et al., 2022). Proband 1(2II) was 76 years old male without any known HCM gene mutations. The other family members were healthy with a mean age of 62.6 years (range: 49–73 years). The Proband’s mother died of sudden cardiac death; however, his father’s clinical records were unavailable (Figure 2A).

**Figure 2:**
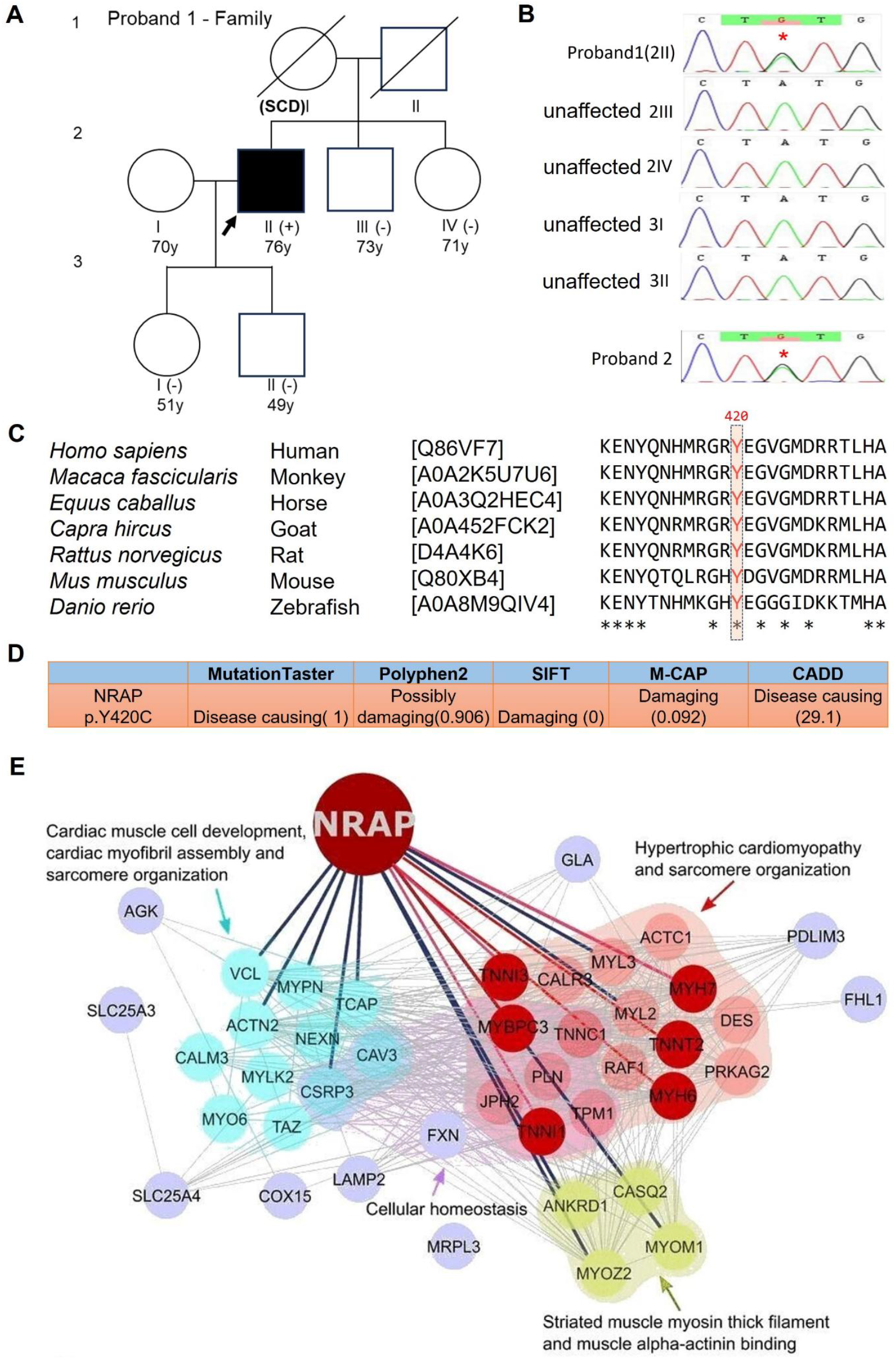
*NRAP* variants in elderly HCM. **(A)** Pedigree of the HCM family, square denotes male and circle denotes female subjects, slash on the symbol represents members who had died. SCD-sudden cardiac death. Black color with an arrow represents the proband, (+) represents the presence of NRAP p.Y420C, and (-) represents the absence of the amino acid change. **(B)** Sanger sequencing chromatograms showing *NRAP* variant (c.1259A>G, p.Y420C) in the Proband 1 (2II) and unrelated HCM patient (Proband 2) and WT sequence in his family members. **(C)** Protein sequence alignment of NRAP across different species showing the conserveness of the Y420 amino acid. **(D)** Pathogenicity prediction scores using in silico tools showing the pathogenic nature of p.Y420C substitution. **(E)** PPI network analysis of NRAP with HCM associated genes and pathway enrichment analysis.

Segregation analysis also revealed that this variant was absent in the remaining unaffected family members (Figure 2B). Notably, the NRAP p.Y420 residue is highly conserved among different species and has been predicted to be pathological using various *in silico* computational tools (Figure 2C and D). The *NRAP* variant was absent in the exomes of Indian controls (n=1029)(A. Jain et al., 2021) and Indian Americans (n=173). Furthermore, it has not been reported in disease databases, including GenoM_2_P, TOPMED, and ClinVar. However, it was present in gnomAD v4.0, with an ultra-rare frequency (MAF= 0.0002%, 3 in 1461686 alleles).

We did not observe the *NRAP* variant in the Indian Wellderly dataset (i-DHANS), strengthening its disease-causing nature. We screened for the *NRAP* variant in two additional elderly control cohorts, including Indian-specific centenarians who were not wellderly individuals (n=93) and Caucasian-specific wellderly individuals (n=500) (Erikson et al., 2016; A. Jain et al., 2021). Notably, *NRAP* p.Y420C was absent in all these exomes. These data provide strong evidence supporting the pathological role of the *NRAP* variant in elderly patients with HCM. Next, we analyzed unrelated patients with HCM (n=335) using targeted resequencing of the respective *NRAP* variants in a replication cohort. The baseline characteristics of the study cohort are provided in Supplementary Table 1. Notably, we identified the same *NRAP* variant in an additional patient with HCM (Proband 2) (Figure 2B).

Genotype-phenotype correlations revealed that Proband 1 (2II) was diagnosed with HCM at the age of 61 years (onset age). The patient had LVIDd and LVIDs measurements of 44 mm and 24 mm, respectively, with an LVEF of 68% and an IVS of 22 mm. Another patient with HCM (Proband 2) from the replication cohort, who experienced onset at the age of 64 years, showed the following echo parameters: LVIDd, 43 mm; LVIDs, 29 mm; LVEF, 60%; and IVS, 21 mm.

### 3.3 Protein-Protein Interaction (PPI) network analysis

Next, we investigated the potential molecular link between NRAP and HCM by conducting a Protein-protein interaction (PPI) network analysis between NRAP and known HCM causing gene encoding proteins. The results of network analysis showed that NRAP was strongly associated with several HCM-associated proteins, including six elderly and late-onset HCM gene-encoding proteins, as depicted in Figure 2E. Furthermore, pathway enrichment analysis indicated a significant association between NRAP and HCM pathways, such as sarcomere organization, cardiac muscle cell development, cardiac myofibril assembly, striated muscle myosin thick filaments, muscle alpha-actinin binding, and cellular homeostasis (Figure 2E).

### 3.4 Functional implications of the NRAP variant

To understand the functional consequences of *NRAP* variant (p.Y420C) in relation to its interacting proteins, we performed immunoprecipitation experiments in cells expressing Kelch-like protein 41 (KLHL41) or α-actinin (Carroll et al., 2001; Lu et al., 2003) with either NRAP wild-type or p.Y420C. The Y420 residue is situated in the simple repeat domain of NRAP, which is known to interact with HCM-associated proteins such as α-actinin and KLHL41 (Figure 3A). The results of our immunoprecipitation experiments indicated that the p.Y420C variant of NRAP exhibited diminished interactions with KLHL41 in comparison to the wild-type NRAP (Figure 3B-D). Additionally, the p.Y420C variant of NRAP displayed reduced interactions with its other interacting partner, α-actinin, in comparison to the wild-type (Figure 3E-G). These results indicate that NRAP p.Y420C contributes to cardiomyopathy in the elderly through a mechanism that involves loss of binding to α-actinin/KLHL41.

**Figure 3:**
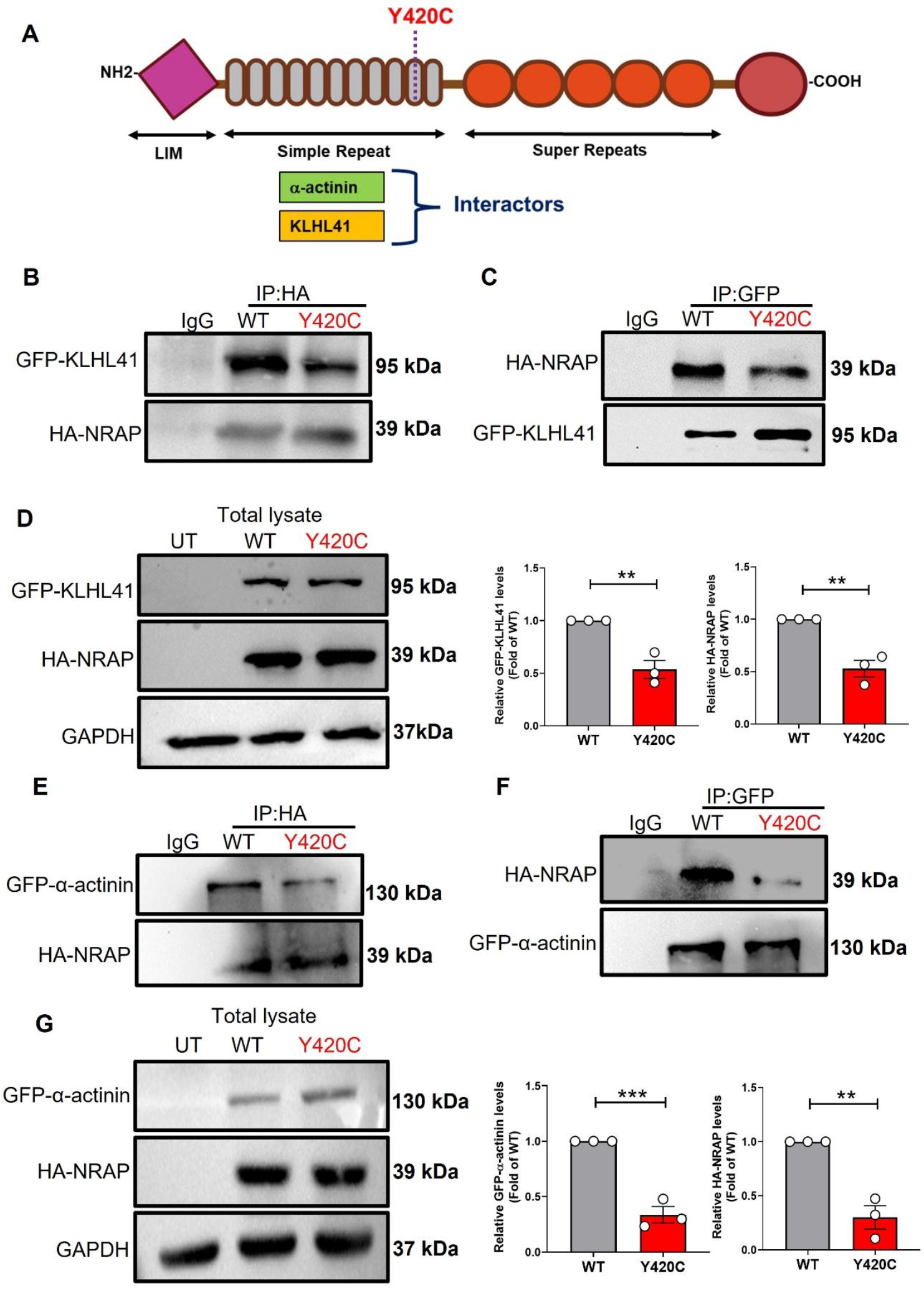
NRAP p.Y420C displays reduced interaction with KLHL41 and α-actinin. **(A)** Schematic representation of the domains of NRAP with its two interactors (α-actinin and KLHL41) along with the location of the mutation site. HEK293T cells were transiently transfected with HA-NRAP (WT/Y420C) and GFP-KLHL41 or GFP-α-actinin. The cell lysates were immunoprecipitated (IP) with anti-HA and anti-GFP antibodies followed by immunoblotting analysis with indicated antibodies. **(B)** Representative immunoblots for HA-NRAP pulldown. **(C)** Representative immunoblots for GFP- KLHL41 pulldown. **(D)** Representative immunoblots for the total lysates including untransfected cell lysate control (UT) with indicated antibodies. **(E)** Representative immunoblots for HA-NRAP pulldown. **(F)** Representative immunoblots for GFP-α-actinin pulldown. **(G)** Representative immunoblots for the total lysates including untransfected cell lysate control (UT) with indicated antibodies. Densitometric analysis for the immunoblots B&C and E&F normalized to WT, respectively. Values are shown as means ± SEM with each experiment performed in triplicate (n=3). Significance was evaluated by unpaired two-tailed Student’s t test **p<0.01 and ***p<0.001.

## 4. Discussion

Here, we sequenced and generated a new Indian wellderly dataset (i-DHANS), which facilitates the identification of *NRAP* as an elderly HCM gene. NRAP is crucial for cardiac muscle thin filament assembly, actin cytoskeleton organization, and myofibril assembly in cardiomyocytes(Carroll et al. 2001). NRAP is a muscle-specific protein with three domains: the LIM domain (LIM), nebulin-related simple repeats (NSR), and super repeats homologous to nebulin (SR)(Luo et al., 1997). During myofibril assembly, NRAP binds to the membrane, facilitating cardiac muscle contraction and relaxation via complexes containing β-integrins, talin, and vinculin. The mutated residue that we observed in the patients (p.Y420) is located in the nebulin-related simple repeat domain (NSR), which contains binding sites for α-actinin and KLHL41 (Figure 3A)(Carroll et al., 2001; Lu et al., 2003). Previous studies have shown that NSR repeats, along with the LIM and SR of NRAP, are critical for the pathogenesis of different types of cardiomyopathies. For example, compound mutations in NRAP, including frameshift mutations in the NSR and SR domains, cause loss of function and result in pediatric dilated cardiomyopathy (PDCoM)(Vasilescu et al., 2018). Furthermore, compound heterozygous missense mutations in the NSR domain have been identified in patients with myofibril myopathy (MFM)(D’Avila et al. 2016). In line with these data, a recent study showed that pathologically increased levels of NRAP contribute to muscle dysfunction in nemaline myopathy (NM) by interacting with KLHL41. NM is strongly associated with HCM(Jirka et al., 2019; Kim et al., 2011; Mir et al., 2012).

Our results suggest that NRAP interactions with α-actinin and KLHL41 are important in pathological processes. NRAP is known to help in the assembly of α-actinin on the z-disc, where it helps maintain the structure of the sarcomere in various models(Manisastry et al., 2009). On the other hand, KLHL41 is a CUL3 E3 ligase adaptor that targets NRAP for degradation by the proteasome, affecting NRAP homeostasis in the system(Jirka et al., 2019). A knockout mice of KLHL41 exhibit severe NM with increased NRAP levels (Ramirez-Martinez et al., 2017). As NRAP plays a crucial role in the regulation of sarcomere formation, its dysregulation affects cardiomyocyte homeostasis, leading to HCM. In line with this, through our co-immunoprecipitation studies, we observed a significant loss of interaction between NRAP p.Y420C and α-actinin or KLHL41 compared to WT NRAP, suggesting that these interactions are affected. As direct evidence, we and others showed that a similar kind of loss-of-function effect was observed in patient cardiac tissue samples with *NRAP* variants (Lesurf et al., 2022; Truszkowska et al., 2017).

To date, all documented *NRAP* variants have been identified as truncating mutations linked to early onset diseases(Koskenvuo et al., 2021; Vasilescu et al., 2018). In contrast, our study suggests that a missense amino acid change at the 420^th^ residue of NRAP is strongly associated with elderly HCM. The NRAP p.Y420C mutant showed significantly less interactions with its interacting partners such as KLHL41 and α-actinin suggesting a potential loss of function mechanism.

## 5. Conclusion

Taken together, we generated the first Indian Wellderly healthy control dataset (i-DHANS) enriched with ethnic-specific genetic factors against age-related diseases. Using the i-DHANS database and the Indian HCM cohort, we provide evidence that *NRAP* is associated with elderly HCM through a loss of function mechanism. Additionally, Indian Wellderly dataset provides a valuable resource for identifying disease genes associated with various conditions.

## Data availability

The data that support the findings of this study are available from the corresponding author upon reasonable request.

## Acknowledgements

We thank all the Wellderly controls, patients and families for the participation in this study. We also thank Ms. Paulami Dey for her kind help in Sanger sequencing.

## Contributions

PSD conceived, designed, analyzed and drafted the manuscript. PC performed and analyzed the functional studies. SA designed and performed exome analysis and i-DHANS dataset. RA helped in functional analysis. VJR helped in i-DHANS dataset. RK, RR, KSM, JS, SS, SB, HA provided Samples for the study. PC and SA performed data analysis and helped in drafting the manuscript. All authors read and approved the final manuscript.

## Consent to participate

Informed written consent was obtained from participating individuals

## Ethics approval

This study was performed in line with the principles of the Declaration of Helsinki. Approval was granted by the Ethics Committee of Institution (Reference number: inStem/IEC-10/001).

## Supplementary information

Supplementary table 1: The details of the baseline characteristics of the HCM patients.

Supplementary table 2: Validation and our classification of HCM associated VUS or conflicting interpretation variants for their pathogenicity in i-DHANS database.

