## Supplementary table 1 for "Indian Wellderly exomes underscore *NRAP* roles in elderly hypertrophic cardiomyopathy"

**Supplementary table 1:** The details of the baseline characteristics of the HCM patients

| ***Variable*** | ***HCM Cohort* (n=335)** |
|---|---|
| ***Demographics*** | |
| Age, year (mean ± SD) | 44.07±12.77 |
| Male, n (%) | 218 (65.07%) |
| Family history of HCM, n (%) | 100 (29.85%) |
| Competitive sport, n (%) | 1 (0.30%) |
| Chest pain, n (%) | 214 (63.88%) |
| Dyspnea, n (%) | 148 (44.18%) |
| Syncope, n (%) | 194 (57.91%) |
| ***Comorbidities*** | |
| CAD, n (%) | 1 (0.30%) |
| Alcohol | None |
| Smoking | None |
| HTN, n (%) | 5 (1.49%) |
| ***ECG data*** | |
| Atrial fibrillation, n (%) | 67 (20%) |
| LBBB, n (%) | 14 (4.18%) |
| RBBB, n (%) | 14 (4.18%) |
| AV block, n (%) | 7 (2.09%) |
| T-wave abnormality, n (%) | 251 (74.93%) |
| ***Echocardiography data*** | |
| LVIDd, mm | 43.63±5.38 |
| LVIDs, mm | 28.15±5.25 |
| LVOTO, n (%) | 117 (34.93%) |
| IVSd, mm | 19.51±10.99 |
| LVEF (%) | 65.56±5.75 |
| ***Septal morphology*** | |
| Asymmetric, n (%) | 273 (81.49%) |
| Apical, n (%) | 30 (8.96%) |
| Concentric, n (%) | 32 (9.55%) |
| ***Cardiac MRI (n=82)*** | |
| Presence of LGE, n (%) | 57 (69.51%) |
| ***Intervention/Medication*** | |
| Myectomy, n (%) | 3 (0.90%) |
| Sudden cardiac death, n (%) | 14 (4.18%) |
