## Supplementary table 2 for "Indian Wellderly exomes underscore *NRAP* roles in elderly hypertrophic cardiomyopathy"

**Supplementary table 2:** Validation and our classification of HCM associated VUS or conflicting interpretation variants for their pathogenicity in i-DHANS database.

| Gene variant | gnomAD v4 | gnomAD classification | i-DHANS (Indian Wellderly database) | Our classification |
|---|---|---|---|---|
| MYH7 p.D809Y | Present | VUS[#] | Present | PNDC* |
| MYH6 p.E1829Q | Present | VUS | Present | PNDC |
| ALPK3 p.T82M | Present | Conflicting interpretation | Present | PNDC |
| MYL2 p.F103C | Present | Conflicting interpretation | Present | PNDC |
| FLNC p.T1317A | Present | Conflicting interpretation | Present | PNDC |
| CSRP3 p.G6R | Present | Conflicting interpretation | Present | PNDC |
| JPH2 p.R572H | Present | VUS | Present | PNDC |

*PNDC: Probably non disease causing

[#]VUS: Variant of uncertain significance
